# A functional overlap between actively transcribed genes and chromatin boundary elements

**DOI:** 10.1101/2020.07.01.182089

**Authors:** Caroline L Harrold, Matthew E Gosden, Lars L P Hanssen, Rosa J Stolper, Damien J Downes, Jelena M. Telenius, Daniel Biggs, Chris Preece, Samy Alghadban, Jacqueline A Sharpe, Benjamin Davies, Jacqueline A. Sloane-Stanley, Mira T Kassouf, Jim R Hughes, Douglas R Higgs

## Abstract

Mammalian genomes are subdivided into large (50-2000 kb) regions of chromatin referred to as Topologically Associating Domains (TADs or sub-TADs). Chromatin within an individual TAD contacts itself more frequently than with regions in surrounding TADs thereby directing enhancer-promoter interactions. In many cases, the borders of TADs are defined by convergently orientated boundary elements associated with CCCTC-binding factor (CTCF), which stabilises the cohesin complex on chromatin and prevents its translocation. This delimits chromatin loop extrusion which is thought to underlie the formation of TADs. However, not all CTCF-bound sites act as boundaries and, importantly, not all TADs are flanked by convergent CTCF sites. Here, we examined the CTCF binding sites within a ∼70 kb sub-TAD containing the duplicated mouse α-like globin genes and their five enhancers (5’-R1-R2-R3-Rm-R4-α1-α2-3’). The 5’ border of this sub-TAD is defined by a pair of CTCF sites. Surprisingly, we show that deletion of the CTCF binding sites within and downstream of the α-globin locus leaves the sub-TAD largely intact. The predominant 3’ border of the sub-TAD is defined by a steep reduction in contacts: this corresponds to the transcribed α2-globin gene rather than the CTCF sites at the 3’-end of the sub-TAD. Of interest, the almost identical α1- and α2-globin genes interact differently with the enhancers, resulting in preferential expression of the proximal α1-globin gene which behaves as a partial boundary between the enhancers and the distal α2-globin gene. Together, these observations provide direct evidence that actively transcribed genes can behave as boundary elements.

**Significance Statement:** Mammalian genomes are complex, organised 3D structures, partitioned into Topologically Associating Domains (TADs): chromatin regions that preferentially self-interact. These chromatin interactions are thought to be driven by a mechanism that continuously extrudes chromatin loops, forming structures delimited by chromatin boundary elements and reflecting the activity of enhancers and promoters. Boundary elements bind architectural proteins such as CCCTC-binding factor (CTCF). Previously, an overlap between the functional roles of enhancers and promoters has been shown. However, whether there is overlap between enhancers/promoters and boundary elements is not known. Here, we show that actively transcribed genes can also behave as boundary elements, similar to CTCF boundaries. In both cases, multi-protein complexes bound to these regions may stall the process of chromatin loop extrusion.

## Introduction

Gene expression throughout development and differentiation is controlled by an interplay between three fundamental *cis*-acting regulatory elements: enhancers, promoters, and boundary elements. Although each type of element is classified by a working definition which enables researchers to establish the syntax of the genome, it is becoming increasingly clear that there is some overlap in the functional roles of these elements as currently defined. For example, some enhancers act as promoters (1-4) and some promoters may also act as enhancers (2, 5-7). Whether enhancers and promoters can also act as boundary elements has been less well studied.

In mammals, boundary elements are frequently located at the borders of large (∼50-2000 kb) regions of chromatin referred to as Topologically Associating Domains (TADs or sub-TADs; self-interacting domains that are nested within larger TADs with a median size of 185 kb) (8-11). TADs are defined as regions of self-interacting chromatin, in that chromatin within a TAD has a higher contact frequency with itself than with regions in surrounding TADs (8, 9, 12). This is thought to ensure that enhancers predominantly interact with promoters present in the same TAD, adding to the specificity of gene regulation. Current models propose that TADs are formed by the extrusion of chromatin loops via translocation of the cohesin complex (13-15). Importantly, boundary elements recruit the zinc finger CCCTC-binding factor (CTCF) which interrupts the translocation of cohesin in an orientation dependent manner and stabilises this protein complex on chromatin. Consistent with this model, cohesin has been shown to be enriched at active boundary elements (16-20). Deletion or inversion of boundary elements often alters the extent of self-interacting TADs and may enable the formation of new enhancer-promoter contacts often producing aberrant gene regulation (9, 15, 21-30).

Despite this coherent model integrating the role of enhancers, promoters, and boundary elements which relates genome structure to gene expression, not all CTCF-bound sites act as boundaries (8) and, importantly, not all TADs are flanked by convergent CTCF sites (11, 24). For example, deletion of a CTCF-rich region in the *Firre* locus found that its TAD boundary is preserved, providing evidence for CTCF-independent boundaries (31). Global depletion of CTCF results in a loss (32, 33) or weakening (34) of ∼80% of TADs across the genome; but not all TADs depend on CTCF. Of interest, removal of CTCF does not lead to widespread mis-regulation of gene expression (32, 33). Together, these observations suggest that elements other than CTCF binding sites might act as functional boundaries. Previous reports have proposed that actively transcribed genes may play such a role. First, Transcriptional Start Sites (TSSs) of housekeeping genes are enriched at TAD borders (8, 35) and, second, the act of transcription can affect 3D genome structure independently of CTCF (36-38). However, whether an actively transcribed gene can behave as a boundary, in a similar manner to a CTCF element, has not been previously tested in mammalian systems.

The duplicated mouse α-like globin genes (*Hba-α1* and *Hba-α2*) and their five enhancers (R1, R2, R3, Rm, and R4) form a very well-characterised, small ∼70 kb tissue-specific sub-TAD in erythroid cells, arranged 5’-R1-R2-R3-Rm-R4-*Hba-α1*-*Hba-α2*-3’. In the past, this locus has been extensively used to establish the principles underpinning mammalian gene regulation and relating genome structure to function (26, 39-44). The α-globin sub-TAD is flanked by several, largely convergent CTCF binding sites (26). We have previously shown that *in vivo* deletion of two CTCF binding sites at the upstream border of the α-globin locus results in an expansion of the sub-TAD and the incorporation of three upstream genes into a newly formed sub-TAD. These three genes become upregulated in erythroid cells via interactions with the α-globin enhancers (26). Therefore, the intact 5’ boundary normally delimits enhancer interactions and thereby contributes to tissue-specific regulation of gene regulation. However, it is not known which, if any, of the regulatory elements produce a similar boundary at the 3’ limit of the α-globin sub-TAD.

To investigate the boundary elements within and downstream of the sub-TAD, we used CRISPR-Cas9 mediated targeting to generate mouse models with mutations of relevant CTCF binding sites. Specific CTCF sites were targeted individually and in informative combinations. We found that the 3’ border of the TAD is only minimally affected by inactivation of any of the tested CTCF sites either individually or in combination. Rather, this border is predominantly defined by the actively transcribed downstream α2-globin gene (*Ηba-α2*). In addition, we found that, when transcribed, the upstream α1-globin gene (*Hba-α1*) acts as a partial barrier to enhancer-promoter interactions with the downstream α2-globin gene. Together our findings demonstrate that actively transcribed genes themselves may behave as boundary elements.

## Results

### Deletion of downstream CTCF sites results in only minor expansion of the self-interacting α-globin sub-TAD with no changes in gene expression

We have previously characterised the sequence and orientation of 16 CTCF binding sites within and flanking the mouse α-globin locus (26). In general, the sub-TAD is flanked by convergently orientated sites (Figure 1). Using NG Capture-C (hereafter referred to as Capture-C) in erythroid cells, we have previously shown and confirm here (Figure 1) that two CTCF binding sites (HS+44/+48) at the 3’ end of the α-globin locus display diffuse, weak interactions with the two CTCF-bound sites (HS-38/-39) that constitute the upstream (5’) boundary of the sub-TAD (26). These CTCF sites do not interact at all with the active enhancer elements (R1-R4 & Rm) within the α-globin sub-TAD. Therefore, we initially considered that HS+44/+48 might delimit the interactions of the α-globin enhancers with promoters lying downstream of the sub-TAD in a similar way to that of HS-38/-39 at the upstream border. We therefore used CRISPR-Cas9 mediated mutagenesis to generate mice with deletions in the binding sequences of these two CTCF sites (Δ44-48; Figure 1 and Supplementary Figure 1a).

**Figure 1:**
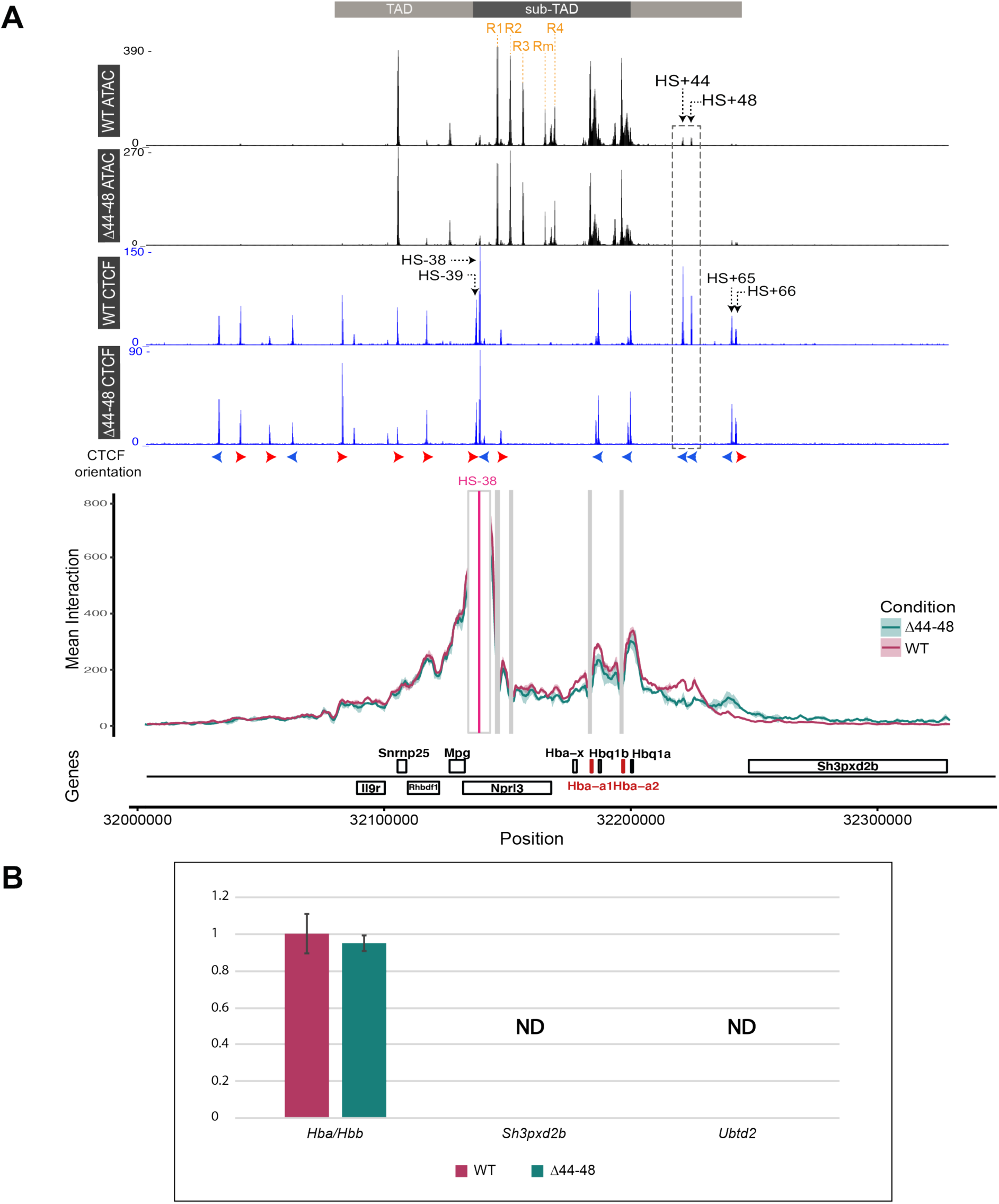
Characterisation of the deletion of CTCF-bound sites downstream of the α-globin locus. **A**: Top tracks show profiles for ATAC-seq and CTCF ChIP-seq in primary erythroid cells isolated from WT (26) and Δ44-48 mice (Ter119+) for the α-globin locus on chromosome 11. Profiles show normalised (RPKM) and averaged data from three biological replicates. The individual α-globin enhancer elements are highlighted in orange (R1-R4 & Rm). The horizontal grey bars above the tracks represent the ∼70 kb α-globin sub-TAD (dark grey) nested within a larger ∼165 kb TAD (light grey). The orientation of CTCF motifs is shown under peaks by red (forward) and blue (reverse) arrows. NG Capture-C interaction profiles of the α-globin locus from the viewpoint of HS-38 (pink), with a 1 kb exclusion zone around the viewpoint, in WT (red) and Δ44-48 (green) Ter119+ primary erythroid cells. The profiles represent normalised and averaged unique interactions from three biological replicates, with the halo representing the standard deviation of a sliding 3 kb window. Vertical grey bars denote other capture points included in this experiment. Genes and genomic position below interaction profiles, with positioning of genes above or below the line representing sense (above) and antisense (below) transcription. The α-globin genes are highlighted in red. **B**: Reverse transcription qPCR expression analysis of α- and β-globin mRNA ratio, *Sh3pxd2b* mRNA, and *Ubtd2* mRNA in WT (red) and Δ44-48 (green) Ter119+ erythroid cells, normalised to 18S RNA. Mean and standard deviation of three biological replicates shown. Data normalised to WT.

In erythroid cells, isolated from the spleens of homozygous Δ44-48 mice, mutations of the HS+44/+48 binding sequences resulted in a complete loss of CTCF binding at these sites without affecting CTCF binding to other, nearby sites (Figure 1a). In addition, other than the loss of peaks at the deleted CTCF sites, the chromatin accessibility around the α-globin locus in Δ44-48 erythroid cells remained unaltered when compared to wild-type (WT) erythroid cells, indicating that other tissue-specific regulatory elements remained intact and unaltered. To investigate whether the removal of the HS+44/+48 sites resulted in changes in local genome topology, we performed Capture-C from viewpoints across the α-globin locus in WT and Δ44-48 primary erythroid cells. Capture-C profiles from the viewpoint of the functional upstream boundary (CTCF site HS-38) show that in absence of CTCF binding to the HS+44/+48 sites, the diffuse interactions over the downstream sites had shifted to the next pair of downstream CTCF binding sites (HS+65/+66), resulting in a minor expansion of the sub-TAD (Figure 1a). Moreover, Capture-C profiles from the viewpoint of the R1 enhancer showed that this shift was accompanied by only slightly increased interactions between R1 and the region of chromatin downstream of the deleted HS+44/+48 sites. This region does not contain any regulatory elements, showing that this expansion occurs even without the formation of new interactions between defined *cis*-regulatory elements (Supplementary Figure 2). Importantly, the altered local chromatin interactions in Δ44-48 erythroid cells were not accompanied by any changes in local gene expression. The two genes directly downstream of the α-globin locus (*Sh3pxd2b* and *Ubtd2*) are not expressed in WT primary erythroid cells, and RT-qPCR analysis found no detectable difference in expression of *Sh3pxd2b, Ubtd2*, or *Hba-α1/2* in Δ44-48 erythroid cells when compared to WT erythroid cells (Figure 1b).

**Figure 2:**
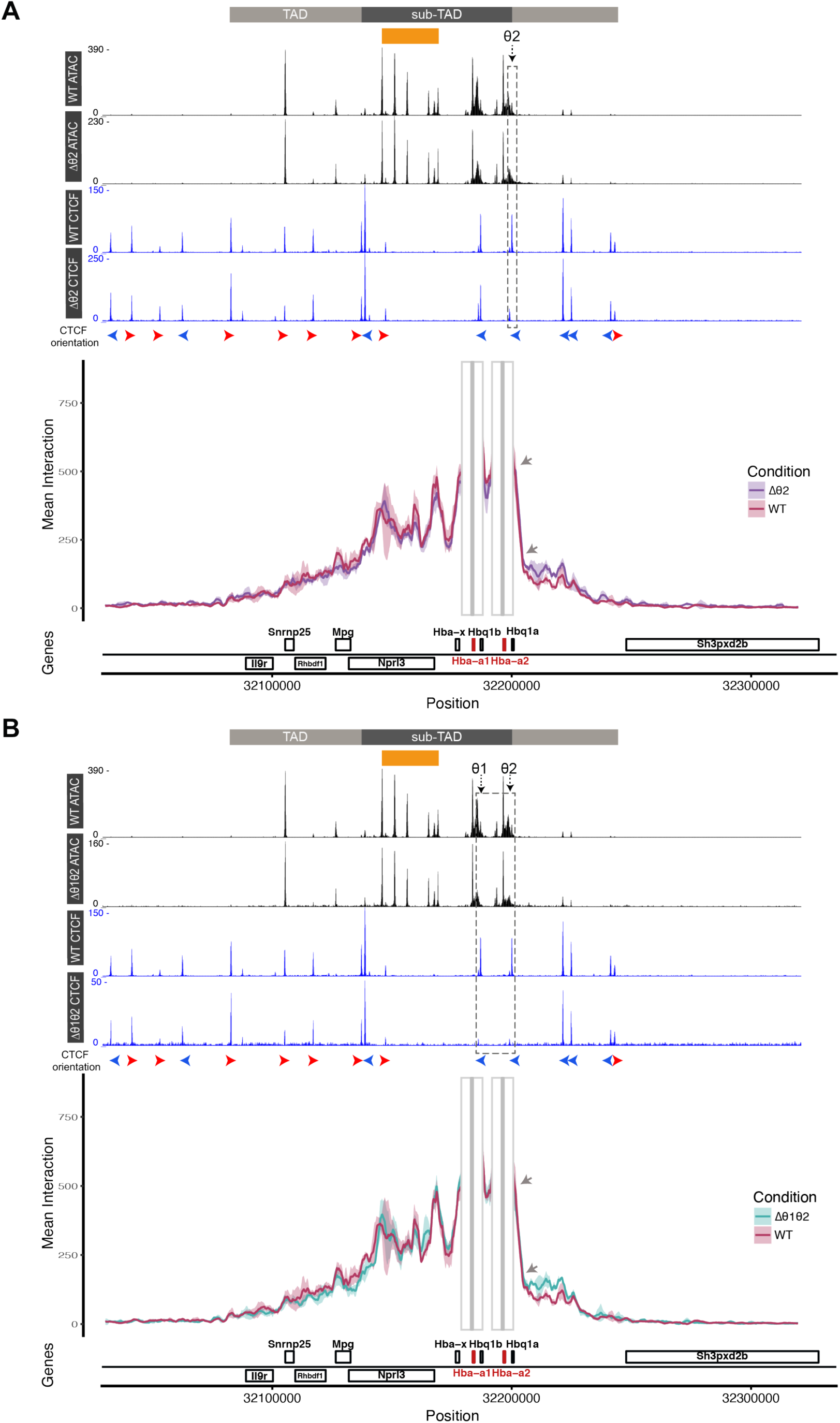
Deletion of CTCF-bound sites leaves the 3’ boundary of the α-globin sub-TAD largely intact. Interaction profiles from the combined viewpoints of the α-globin promoters in Δθ2 (**A**) and Δθ1θ2 (**B**) erythroid cells. In **A, B**: Top tracks show profiles for ATAC-seq and CTCF ChIP-seq in primary erythroid cells isolated from WT (26) and the respective mouse model (Ter119+) for the α-globin locus on chromosome 11. Profiles show normalised (RPKM) and averaged data from three biological replicates. The α-globin enhancer region is represented by an orange box. The horizontal grey bars above the tracks represent the ∼70 kb α-globin sub-TAD (dark grey) nested within a larger ∼165 kb TAD (light grey). The orientation of CTCF motifs is shown under peaks by red (forward) and blue (reverse) arrows. NG Capture-C interaction profiles of the α-globin locus from the viewpoints of the α-globin promoters (grey), with a 1 kb exclusion zone around the viewpoints, in WT (red), Δθ2 (purple), and Δθ1θ2 (teal) Ter119+ primary erythroid cells. The profiles represent normalised and averaged unique interactions from three biological replicates, with the halo representing the standard deviation of a sliding 3kb window. Grey arrows denote the 3’ edge of the α-globin sub-TAD. Genes and genomic position below interaction profiles, with positioning of genes above or below the line representing sense (above) and antisense (below) transcription. The α-globin genes are highlighted in red.

Taken together, these findings show that rather than behaving as a strong boundary element, the HS+44/+48 CTCF sites behave as a minor boundary to loop extrusion and the potential for chromatin interactions. However, unlike the previously characterised 5’ boundary and other boundaries described in the literature (9, 15, 22-24, 26, 27, 29, 30), removal of these CTCF sites does not lead to any changes in gene expression within or flanking the α-globin gene cluster.

### The actively transcribed α2-globin gene acts as the downstream boundary of the α-globin sub-TAD

Since the HS+44/+48 sites do not constitute the 3’ boundary of the α-globin sub-TAD, we next considered the more proximal CTCF sites that coincide with the θ1/2 genes (*Hbq1b* and *Hbq1a*) (Figure 1), which are situated inside the α-globin sub-TAD and within the duplicated region of the α-globin locus. The θ1/2 genes are α-like genes of unknown function (45). In addition to displaying diffuse interactions with the downstream HS+44/+48 sites, the upstream (5’) boundary of the α-globin sub-TAD also diffusely interacts with the θ1/2 CTCF-bound sites (26) (Figure 1). Furthermore, we have previously shown that Capture-C interaction profiles from the viewpoint of any active regulatory element inside the undisturbed sub-TAD display a pronounced reduction in interactions immediately downstream of the 3’ α2-globin gene (*Hba-α2*), which, within the resolution of these studies, appears to coincide with the θ2 CTCF binding site (26, 39-41). Therefore, it seemed possible that this CTCF-bound site could act as the 3’ boundary of the α-globin sub-TAD. To investigate the roles of CTCF sites inside an active sub-TAD with respect to genome structure and gene regulation, we used CRIPSR-Cas9 mediated mutagenesis to generate mice with deletions at the θ CTCF sites individually or in combination (Δθ1, Δθ2, and Δθ1θ2; Figure 2, Figure 3, Supplementary Figure 1b, and Supplementary Figure 3).

**Figure 3:**
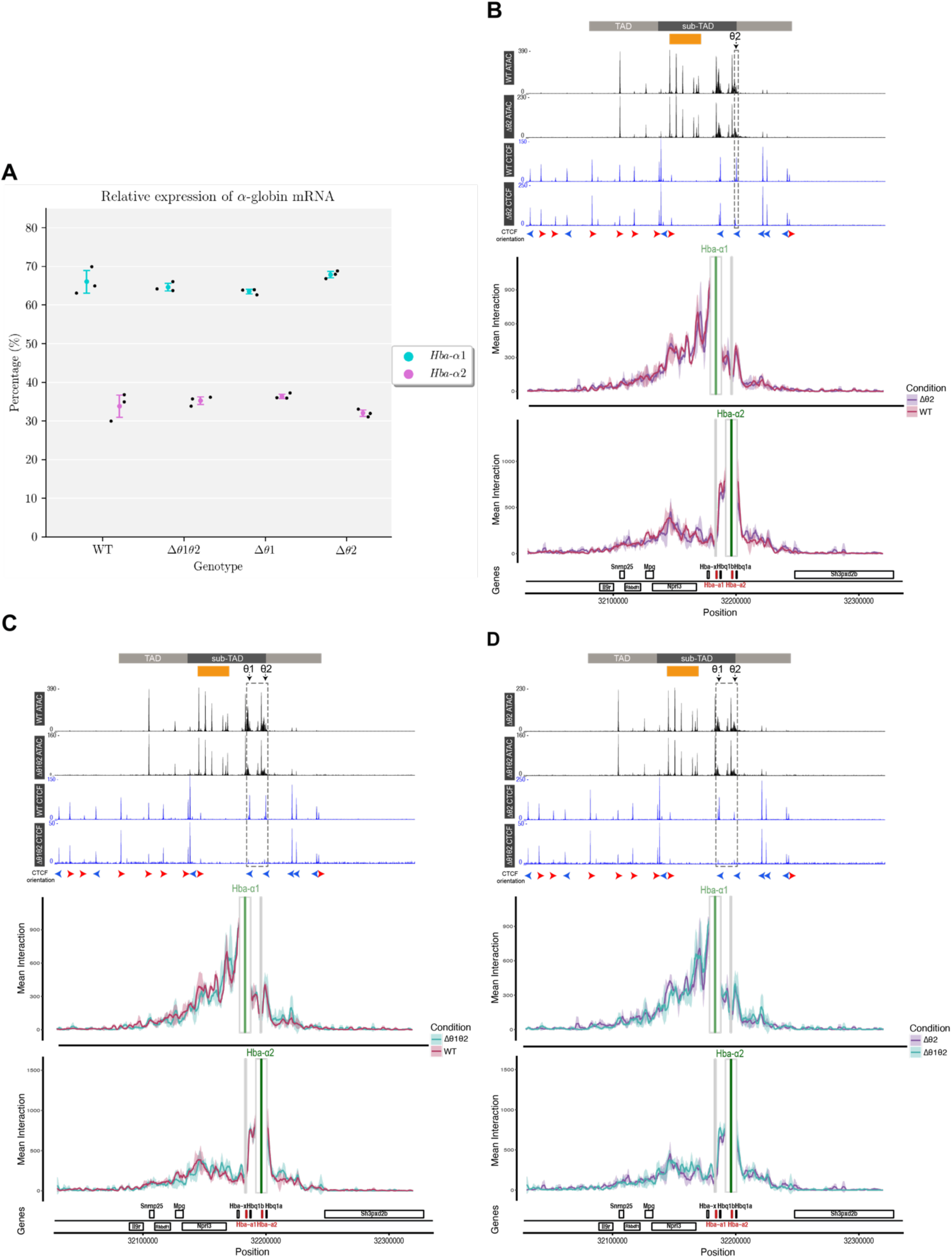
CTCF does not regulate the differential interactions and expression of the α-globin genes. **A**: Relative expression of *Hba-α1/2* mRNA in WT, Δθ1θ2, Δθ1, and Δθ2 primary erythroid cells (Ter119+). Variant calling analysis performed on Poly(A)+ RNA-seq data from biological triplicates revealed percentage of reads originating from transcripts of *Hba-α1* (teal) or *Hba-α2* (pink). Mean and standard deviation for each model shown, and each point represents a biological replicate. **B**: Effects of the deletion of θ2 on local chromatin accessibility, CTCF binding, and *Hba-α1/2*-specific interaction profiles. **C**: Effects of the combined deletion of θ1and θ2 on local chromatin accessibility, CTCF binding, and *Hba-α1/2-*specific interaction profiles. **D**: Comparison of *Hba-α1/*2-specific interaction profiles in Δθ2 and Δθ1θ2 erythroid cells. In **A, B, C**: Top tracks show profiles for ATAC-seq and CTCF ChIP-seq in primary erythroid cells isolated from WT (26) and the respective mouse model (Ter119+) for the α-globin locus on chromosome 11. Profiles show normalised (RPKM) and averaged data from three biological replicates. The α-globin enhancer region is represented by an orange box. The horizontal grey bars above the tracks represent the ∼70 kb α-globin sub-TAD (dark grey) nested within a larger ∼165 kb TAD (light grey). The orientation of CTCF motifs is shown under peaks by red (forward) and blue (reverse) arrows. NG Capture-C interaction profiles of the α-globin locus from the viewpoints of the α-globin promoters (*Hba-α1* in light green; *Hba-α2* in dark green), with a 1 kb exclusion zone around the viewpoints, in WT (red), Δθ2 (purple), and Δθ1θ2 (teal) Ter119+ primary erythroid cells. The profiles represent normalised and averaged unique interactions from three biological replicates, with the halo representing the standard deviation of a sliding 3 kb window. Genes and genomic position below interaction profiles, with positioning of genes above or below the line representing sense (above) and antisense (below) transcription. The α-globin genes are highlighted in red.

We therefore next analysed primary erythroid cells from Δθ1, Δθ2, and Δθ1θ2 homozygous mice and showed that mutations of the CTCF binding sequences resulted in a complete loss of CTCF binding at the specifically targeted sites (Figure 2, Figure 3, and Supplementary Figure 3). Again, there were no additional changes in chromatin accessibility around the α-globin locus in erythroid cells derived from any of the three θ mouse models when compared to WT erythroid cells (Figure 2, Figure 3, and Supplementary Figure 3). To investigate whether removal of CTCF sites within and immediately downstream of the active α-globin locus caused changes to the sub-TAD structure, we performed Capture-C from the viewpoint of the α-globin promoters in WT, Δθ2, and Δθ1θ2 primary erythroid cells. Surprisingly, in both Δθ2 and Δθ1θ2 erythroid cells, the 3’ boundary of the sub-TAD remained largely intact, and the pronounced reduction in chromatin interactions downstream of the α2-globin gene persisted, despite the loss of CTCF binding at θ2 and both θ sites, respectively (Figure 2; grey arrows). In addition, the overall sub-TAD structure remained largely unaffected. Hence, the θ1/2 CTCF binding sites are not essential to form the 3’ boundary of the sub-TAD and these results show that the 3’ boundary of the sub-TAD is marked by the active α2-globin gene itself.

### Investigating the role of CTCF sites lying within the α-globin sub-TAD in regulating gene expression

Although the CTCF sites lying close by and in between the α-globin genes play no role in forming the 3’ boundary of the α-globin sub-TAD, we considered whether they might play a role in fine tuning gene expression within the sub-TAD. In human, the upstream 5’ α-globin gene (*HBA2*; the equivalent of mouse *Hba-α1*) is located closer to the enhancers and is more highly expressed than the downstream 3’ α-globin gene (*HBA1*; the equivalent of mouse *Hba-α2*). *HBA2* produces ∼70% of the total α-globin mRNA (46-48). To determine if this differential expression of the α-globin genes also holds true for mouse, we performed Poly(A)+ RNA-seq on primary WT erythroid cells. The presence of a single SNP in the third exon of *Hba-α1* and *Hba-α2* allows for variant calling analysis of the reads originating from the α-globin transcripts. As in human, *Hba-α1* (the gene closest to the enhancers) accounted for ∼66% of the total α-globin mRNA and *Hba-α2* (the distal gene) accounted for ∼34% (Figure 3a). Consistent with its higher level of expression, we have previously shown that the *Hba-α1* promoter also preferentially interacts with the α-globin enhancers relative to the *Hba-α2* promoter (40). As the θ CTCF sites are situated downstream of each α-globin gene (in the order 5’-α1-θ1-α2-θ2-3’), we investigated whether the θ CTCF sites, and in particular θ1 situated in between the α-globin genes, regulate the differential expression of the mouse α-globin genes and/or their interactions with the α-globin enhancers.

We first performed Poly(A)+ RNA-seq on primary erythroid cells isolated from Δθ1, Δθ2, and Δθ1θ2 mice and used the variant calling analysis described above on the α-globin transcripts. In all three models, the relative proportions of transcripts produced by *Hba-α1* and *Hba-α2* were similar to that of WT (Figure 3a), showing that the loss of CTCF binding between and downstream of the α-globin genes did not affect the preferential expression of *Hba-α1*. Next we investigated whether the loss of CTCF binding around the α-globin genes altered differential interactions of *Hba-α1/2* with the enhancers. From the Capture-C data analysed from the viewpoints of the α-globin promoters in WT, Δθ2, and Δθ1θ2 primary erythroid cells described above, we generated separate interaction profiles for the *Hba-α1* and *Hba-α2* promoters following previously described analysis (40). When erythroid cells from Δθ2 and Δθ1θ2 were analysed, we observed differential interaction profiles of the α-globin promoters as previously reported in WT erythroid cells (Figure 3b,c). To investigate whether the loss of CTCF binding between the α-globin promoters (at θ1) altered the differential interactions, we generated comparisons of the *Hba-α1/2*-specific interaction profiles between Δθ2 and Δθ1θ2 erythroid cells. The only difference between these two models is the mutation at the θ1 site in Δθ1θ2 mice. Again, there were no observable differences between the mutant mouse models (Figure 3d), indicating that the differential interactions of *Hba-α1/2* with the enhancers is not influenced by presence or absence of the θ1 CTCF site.

Therefore, in summary these findings suggest that CTCF binding at θ1 and/or θ2 do not regulate the differential interactions of *Hba-α1/2* with the α-globin enhancers or the preferential expression of the proximal α1-globin gene (*Hba-α1*) compared to the distal α2-globin gene (*Hba-α2*). Rather, it appears that the transcribed α1-globin gene may act as a partial boundary to the α2-globin gene in terms of access to a shared set of enhancers just as the α2-globin gene acts as the downstream boundary of the sub-TAD.

## Discussion

In many ways, the relatively small ∼70 kb sub-TAD containing the mouse α-globin locus is typical of other tissue-specific TADs or sub-TADs seen in mammalian genomes. The cluster of erythroid-specific enhancers, which fulfil the definition of a super-enhancer (41), and the promoters of the α-like genes, are flanked by largely convergent CTCF binding sites which are often considered to act as the structural and functional boundaries of TADs. Current models relating genome structure to function propose that TADs are formed by the extrusion of chromatin loops as the cohesin complex translocates throughout the TAD (13-15). The continuous process of extrusion ultimately brings together all sequences within the self-interacting TAD, including the enhancers and promoters, providing the proximity thought to be required for the activation of transcription. The borders of the TAD are created when cohesin is stalled and stabilised by its interaction with the N-terminal region of CTCF. Our previous studies of the mouse α-globin sub-TAD using chromosome conformation capture (26, 39-44) and super-resolution imaging (39) are entirely consistent with this model involving the interplay between enhancers, promoters, and boundary elements. Here, we have identified the precise elements responsible for forming the boundaries of the sub-TAD and contributing to the differential interactions between the enhancers and the promoters. Surprisingly, we find that the transcriptionally active α-globin genes rather than the anticipated CTCF binding sites play a role as boundary elements within the locus and in creating the downstream boundary of the sub-TAD.

We have previously characterised the upstream 5’ boundary of the α-globin sub-TAD (26) which is marked by two convergent CTCF binding sites (HS-38/-39 in Figure 1). Deletion of these sites leads to extension of the sub-TAD to more distal flanking CTCF site (HS-59) and erythroid-specific activation of three genes incorporated into the extended sub-TAD. Presumably this occurs because the cohesin complex can now translocate beyond these sites. Deletion of two CTCF sites (HS+44/+48) flanking the 3’ end of the α-globin locus which interact with the upstream boundary behaved differently from those at the 5’ boundary. Deletion of these sites led to only a very small increase in the levels of interaction with the downstream flanking region beyond these sites, extending to the next CTCF sites (HS+65/+66). However, there was no associated change in expression of the downstream flanking genes (*Sh3pxd2b* and *Ubtd2*). In effect, removal of the HS+44/+48 sites caused no change in the major transition (grey arrows in Figure 2) between interacting and non-interacting chromatin. This major boundary coincides with the actively transcribed α2-globin gene and/or a CTCF associated with the θ2-globin gene. Removal of this CTCF site did not alter the predominant transition showing that it is the transcribed α-globin gene itself that corresponds to the prominent 3’ boundary of the sub-TAD.

Previous studies have identified between 15,000 and 40,000 CTCF binding sites in the genome (49-53), but despite having common consensus sequences and chromatin signatures, only a small proportion appear to act as boundary elements or to constrain the activity of enhancers (8). A CTCF site lying between the mouse α1- and α2-globin genes appeared to provide an example of an element which might partially block enhancer activity. The identical promoters of the α1- and α2-globin genes interact differently with the enhancers and consequently direct different levels of α-globin mRNA. However, deletion of the internal CTCF binding sites at θ1/2 had no effect on the differential interactions of the two α-globin genes or the preferential expression of the proximal α1-globin gene. This suggests that the α1-globin gene itself acts as a partial boundary between the α2-globin gene and the α-globin enhancers.

Together, our findings on flanking and internal CTCF sites support the proposal that some actively transcribed genes, in a particular context, may themselves behave as boundaries. One mechanism by which this could occur is via competition between promoters and a shared enhancer in a situation where there is unconstrained chromatin looping (Figure 4A). Such a competition model would propose that the promoter of the α1-globin gene outcompetes that of the α2-globin gene for access to the α-globin enhancers in which all enhancer elements appear to act as a single entity (43). However, this seems unlikely since in the context of a freely interacting chromosomal loop the promoters are located at a relatively similar distance from the enhancer region and are identical in sequence. Similar examples of promoter competition have also been proposed in which active promoters are located between an enhancer and another, more distal promoter causing reduced activity of the distal promoter (54-57).

**Figure 4:**
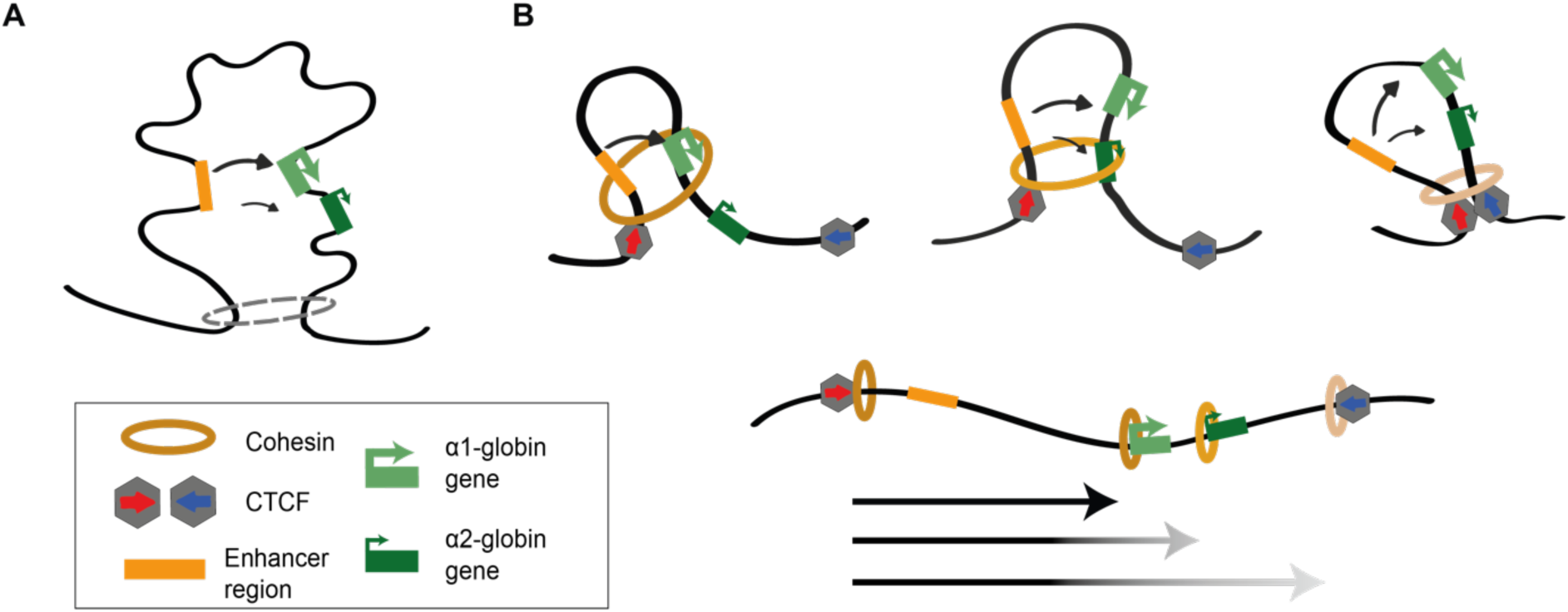
Proposed mechanisms for actively transcribed genes behaving as boundary elements. **A**: Schematic to show that under unconstrained chromatin looping, the α1-globin gene outcompetes the α2-globin gene via promoter competition for access to the shared set of α-globin enhancers. **B**: Schematic to show a dynamic, directional tracking mechanism of chromatin loop extrusion by cohesin from the α-globin enhancers to the promoters. Multi-protein complexes recruited to the actively transcribing genes stall cohesin translocation on chromatin resulting in cohesin retention at active genes, in addition to CTCF binding sites.

An alternative explanation proposes a *directional* tracking mechanism from enhancers to promoters both of which have enriched levels of cohesin. Such a model of directional loop extrusion has been proposed to explain interacting stripes seen in Hi-C maps (58). In this case the anchor of the extruding loop would correspond to the HS-38/-39 sites. Multi-protein complexes recruited to an actively transcribing gene might act, like a CTCF boundary, to stall translocation of the cohesin complex and thereby reduce access of the enhancer to a more distal promoter (Figure 4B). This interpretation of the mouse data presented here, is supported by observations of the orthologous human and sheep α-globin clusters in which, the proximal duplicated α-globin gene is also expressed in preference (∼70 %) to the more distal gene (∼30%) (46-48, 59). In the human there is no CTCF binding site between the α-globin genes. Of relevance, in both human and sheep with further tandem duplications producing chromosomes with two (αα/), three (ααα/), four (αααα/), or five (ααααα/) almost identical α-globin genes (59-62), each additional gene provides a smaller contribution to α-globin mRNA and protein, as in a gradient. Furthermore, in humans, a deletion of the proximal α2-globin gene (-α/) increases the output of the distal α1-globin gene from 20% to 50% (46). By contrast, when there is an inactivating coding mutation in the proximal α2-globin gene (α^M^α/), leaving its promoter and transcription intact, RNA expression from the distal α1-globin gene remains at 20% leading to the severe phenotype seen in patients with such nondeletional mutations (63). A similar situation with a gradient in expression has also been observed for the almost identical duplicated (γγ/), triplicated (γγγ/), and quadruplicated (γγγγ/) γ-globin genes (64). The ratio of expression of the duplicated proximal to the distal γ-globin gene with respect to the β-globin enhancers when they are active in fetal life is again ∼70% to ∼30% (65).

In this model highly transcribed genes might form a barrier to loop extrusion (66), due to accumulation of large amounts of transcriptional machinery and regulatory factors. In these scenarios, cohesin may be prevented from extruding chromatin loops due to the size of multi-protein complexes which may be acting like ‘roadblocks’ via a passive blocking mechanism (Figure 4B). Evidence that supports this comes from structural studies looking at the interaction between CTCF and cohesin. The N terminus of CTCF structurally interacts with cohesin, however, it appears its role is to stabilise cohesin on chromatin; when the N terminus is mutated, cohesin still accumulates at CTCF sites but at lower levels compared to WT (67). This suggests that CTCF is sufficient to block cohesin without a specific interaction and the same could be true for other large proteins.

Therefore, there may be different methods to block loop extrusion; CTCF can directly interact with cohesin causing it to be retained at CTCF-bound sites; however, passive blocking of cohesin by large multi-protein complexes may also occur and could provide a mechanism for how actively transcribed genes can behave as boundaries.

In summary, we provide evidence that in addition to CTCF binding sites, actively transcribed genes may also behave as boundaries in agreement with studies that found that some TAD boundaries are enriched for the promoters of actively transcribed genes, such as housekeeping genes (8, 35). This suggests that an active promoter may have multiple roles in shaping the genome.

## Methods

### Animal procedure

The mutant and wild-type mouse strains reported in this study were generated and maintained on a C57BL/6J background in accordance with the European Union Directive 2010/63/EU and/or the UK Animal (Scientific Procedures) Act 1986, with procedures reviewed by the clinical medicine Animal Welfare and Ethical Review Body (AWERB). Experimental procedures were conducted under project licences PPL 30/3339 and PAA2AAE49. All animals were housed in Individually Ventilated Cages with enrichment, provided with food and water *ad libitum*, and maintained on a 12 h light: 12 h dark cycle (150-200 lux cool white LED light, measured at the cage floor). Mice were given neutral identifiers and analysed by research technicians unaware of mouse genotype during outcome assessment.

### Isolation of erythroid cells derived from adult mouse spleen

Primary Ter119+ erythroid cells were obtained from the spleens of adult mice that were treated with phenylhydrazine as described previously (68). Spleens were mechanically dissociated into single cells suspensions in cold phosphate-buffered saline (PBS; Gibco: 10010023)/10% fetal bovine serum (FBS; Gibco: 10270106) and passed through a 70 μm filter to remove clumps. Cells were washed with cold PBS/10% FBS and resuspended in 10 μl of cold PBS/10% FBS per 10^6^ cells and stained with a 1/100 dilution of anti-Ter119-PE antibody (Miltenyi Biotec: 130-102-336) at 4 °C for 20 minutes. Stained cells were washed with cold PBS/10% FBS and resuspended in 8 μl of cold PBS/0.5% BSA/2 mM EDTA and 2 μl of anti-PE MACS microbeads (Miltenyi Biotec: 130-048-801) per 10^6^ cells and incubated at 4 °C for 15 minutes. Ter119+ cells were positively selected via MACS lineage selection columns (Miltenyi Biotec: 130-042-401) and processed for downstream applications. Purity of the isolated erythroid cells was routinely verified by Fluorescence-Activated Cell Sorting (FACS).

### Generation of mutant mouse strains

Mouse models harbouring mutations of CTCF binding sites around the mouse α-globin locus were generated using CRISPR-Cas9 mediated genome editing by either targeting mouse embryonic stem cells, which were then used in blastocyst injections, or by direct microinjection of zygotes. Preparation of CRISPR-Cas9 expression constructs for targeting of mouse embryonic stem cells and preparation of CRISPR-Cas9 reagents and ssODN templates, as required, for direct microinjection of zygotes were performed as previously described (26). The 20 nucleotide guide sequences used to direct the Cas9 protein to the target CTCF binding sites and the ssODN donor sequences are shown in Supplementary Table 1.

### ATAC-seq

ATAC-seq was performed on 75,000 Ter119+ cells isolated from phenylhydrazine-treated mouse spleens as previously described (69). ATAC-seq libraries were sequenced on the Illumina Nextseq platform using a 75-cycle paired-end kit (NextSeq 500/550 High Output Kit v2.5: 20024906). Data were analysed using an in-house pipeline (70) which uses Bowtie (71) to map reads to the mm9 mouse genome build. PCR duplicates were removed, and biological replicates were normalised to Reads Per Kilobase per Million (RPKM) mapped reads using deeptools bamCoverage (72). Mitochondrial DNA was excluded from the normalisation. For visualisation, ATAC-seq data were averaged across three biological replicates.

### ChIP-seq

CTCF Chromatin immunoprecipitation (ChIP) was performed on 1 × 10^7^ Ter119+ erythroid cells using a ChIP Assay Kit (Millipore: 17-295) according to the manufacturer’s instructions. Cells were crosslinked by a single 10 min 1% formaldehyde fixation. Chromatin fragmentation was performed with the Bioruptor Pico sonicator (Diagenode) for a total sonication time of 4 min (8 cycles) at 4°C to obtain an average fragment size between 200 and 400 bp. Immunoprecipitation was performed overnight at 4 °C with an anti-CTCF antibody (10 µl 07-729, lot: 2836926; Millipore). Library preparation of immunoprecipitated DNA fragments was performed using NEBNext Ultra II DNA Library Prep Kit for Illumina (New England Biolabs: E7645) according to the manufacturer’s instructions. Libraries were sequenced on the Illumina Nextseq platform using either a 75-cycle paired-end kit (NextSeq 500/550 High Output Kit v2.5: 20024906) or a 300-cycle paired-end kit (NextSeq 500/550 Mid Output Kit v2.5: 20024905). Data were analysed using an in-house pipeline (70) which uses Bowtie (71) to map reads to the mm9 mouse genome build. PCR duplicates were removed, and biological replicates were normalised to RPKM mapped reads using deeptools bamCoverage (72). For visualisation, ChIP-seq data were averaged across three biological replicates.

### NG Capture-C

Next-generation Capture-C was performed as previously described (40). A total of 1-2 × 10^7^ Ter119+ erythroid cells were used per biological replicate. We prepared 3C libraries using the DpnII-restriction enzyme for digestion. We added Illumina TruSeq adaptors using the NEBNext Ultra II DNA Library Prep Kit for Illumina (New England Biolabs: E7645) according to the manufacturer’s instructions, and performed capture enrichment using NimbleGen SeqCap EZ Hybridization and Wash Kit (Roche: 05634261001), NimbleGen SeqCap EZ Accessory Kit v2 (Roche: 07145594001), and previously published custom biotinylated DNA oligonucleotides (R1 and HS-38 viewpoints (26); α-globin promoters viewpoints (40)). NG Capture-C data were analysed using the CaptureCompendium toolkit (73) which uses Bowtie (71) to map reads to the mm9 mouse genome build. *Cis* reporter counts for each sample were normalised to 100,000 reporters for calculation of the mean and standard deviation (three biological replicates). Mean reporter counts were divided into 150 bp bins and smoothed using a 3 kb window.

### RNA expression analysis

Total RNA was isolated from 5 × 10^6^ Ter119+ erythroid cells lysed in TRI reagent (Sigma-Aldrich: T9424) using a Direct-zol RNA MiniPrep kit (Zymo Research: R2050). DNase I treatment was performed on the column as recommended in the manufacturer’s instructions but with an increased incubation of 30 min at room temperature (rather than 15 min). To assess relative changes in gene expression by qPCR, cDNA was synthesised from 1 μg of total RNA usingSuperScriptIIIFirst-StrandSynthesisSuperMixforqRT-PCR(Invitrogen, ThermoFisher: 11752-050) according to the manufacturer’s instructions. The ΔΔCt method was used for relative quantification of RNA abundance using TaqMan Universal PCR Master Mix (Applied Biosystems, ThermoFisher: 4304437) and the following TaqMan probes: Mm00845395_s1 (*Hba-α1/2*), Mm01611268_g1 (*Hbb-b1*), Mm00616672_m1 (*Sh3pxd2b*), Mm00612868_m1 (*Ubtd2*), and Mm04277571_s1 (*Rn18s*). For RNA-seq libraries, 1-2 μg of total RNA was depleted of rRNA and globin mRNA using the Globin-Zero Gold rRNA Removal Kit (Illumina: GZG1224) according to the manufacturer’s instructions. To enrich for mRNA, poly(A)+ RNA was isolated, strand-specific cDNA was synthesised, and the resulting libraries prepared for Illumina sequencing using the NEBNext Poly(A) mRNA Magnetic Isolation Module (New England Biolabs: E7490) and the NEBNext Ultra II Directional RNA Library Prep Kit for Illumina (New England Biolabs: E7760) following the manufacturer’s instructions. Poly(A)+ RNA-seq libraries were sequenced on the Illumina Nextseq platform using a 75-cycle paired-end kit (NextSeq 500/550 High Output Kit v2.5: 20024906). Reads were aligned to the mm9 mouse genome build using STAR (74). To perform variant calling analysis on RNA-seq reads originating from α-globin transcripts an in-house variant-caller tool developed by Jelena Telenius was used, which was based on the samtools1 version of mpileup (75) to count variants. Each sample was aligned twice to the mouse mm9 genome: once with *Hba-α1* masked and once with *Hba-α2* masked. The resulting alignments were used as inputs for the variant-caller which counted mismatches at *Hba-α1/2*. Full documentation for the variant-callertoolcanbefoundhere: http://userweb.molbiol.ox.ac.uk/public/telenius/variantApp/variantApp_JTelenius_GPL3_2019.pdf.

## Data availability

All sequencing data have been submitted to the NCBI Gene Expression Omnibus under accession number GSE153209.

## Acknowledgements

This work was supported by Wellcome (Genomic Medicine and Statistics PhD Programme, reference 109110/Z/15/Z; Chromosome and Developmental Biology PhD Programme, reference 099684/Z/12/Z; Wellcome Trust Strategic Award, reference 106130/Z/14/Z; Wellcome Trust Core Award, reference 203141/Z/16/Z) and the Medical Research Council (MRC Core Funding and Project Grant, reference MR/N00969X/1).

## Author contributions

C.L.H., L.L.P.H., B.D., M.T.K., J.R.H. and D.R.H conceived and designed experiments and coordinated and advised on the project. C.L.H., M.E.G. and R.J.S. performed experiments. C.L.H., D.J.D. and J.M.T performed bioinformatic analyses. D.B., C.P., S.A., J.A.S. and J.A.S.-S. generated essential reagents and carried out mice maintenance. C.L.H. and D.R.H. wrote the manuscript. J.R.H. and D.R.H. supervised works carried out.

## Competing interests

J.R.H is a founder and shareholder of Nucleome Therapeutics.

