## Supplementary Figures 1-3; Supplementary Table 1 for "A functional overlap between actively transcribed genes and chromatin boundary elements"

### Supplementary Figure 1: Mutated sequences of CTCF binding sites at the α globin locus

### A

```
WT HS+44     1  ATTAGAAAGCCAGTGGCGCCACCTTGGGGCCCACTCTCAGCTGTTTATGACTACCGGGCA
Δ44-48       1  ATTAGAAAGCCAGTGGCGCCA---------------------------------------

WT HS+44    61  GGAGGTCTCTGGGGCGCCCCCTGCAGGCCACTATAAGTAGGTGCAAGGCTTTTCCAGTAT
Δ44-48      61  --------------------------------------AGGTGCAAGGCTTTTCCAGTAT
```

```
WT HS+48     1  CCGGAGATATAAGAAGGCGACGAGCACCCCCGTGTGGCGGGACTCGGAACTTCGCAGCTC
Δ44-48       1  CCGGAGATATAAGAAGGCGACGAGCACCCCCGTG------GACTCGGAACTTCGCAGCTC
```

### B

```
WT θ1        1  CCTAGGCATCCTGGAACGATGCAGCGCCCCCTGGCGGCCTCTTGTGTTTCAGGACGTCTT
Δθ1          1  CCTAGGCATCCTGGAACGATGCAGCGGGATCC----GCCTCTTGTGTTTCAGGACGTCTT
```

```
WT θ1        1  CCTAGGCATCCTGGAACGATGCAGCGCCCCCTGGCGGCCTCTTGTGTTTCAGGACGTCTT
Δθ1θ2        1  CCTAGGCATCCTGGAACGATGCAGCGC-----------CTCTTGTGTTTCAGGACGTCTT
```

```
WT θ2        1  CCTAGGCATCCTGGAACGATGCAGCGCCCCCTGGCGGCCTCTTGTGTTTCAGGACATCTT
Δθ2/Δθ1θ2    1  CCTAGGCATCCTGGAACGATGCAGCGGGATCC----GCCTCTTGTGTTTCAGGACATCTT
```

**A**: Alignments show WT sequences of HS+44 and HS+48 CTCF binding sites, with the 20 bp core binding motif highlighted in blue (reverse orientation), and deleted sequences of HS+44 and HS+48 CTCF binding sites in Δ44-48 mice (77 bp and 6 bp deletions respectively).

**B**: Alignments show WT sequences of θ1 and θ2 CTCF binding sites, with the 20 bp core binding motif highlighted in blue (reverse orientation), and mutated sequences of θ1 CTCF binding site in Δθ1 (HDR mutation) and Δθ1θ2 mice (11 bp deletion), and the mutated sequence of θ2 CTCF binding in Δθ2 and Δθ1θ2 mice (HDR mutation in both models).

**Supplementary Figure 2: NG Capture-C interaction profiles of the α-globin locus from the viewpoint of the R1 enhancer in Δ44-48 erythroid cells**

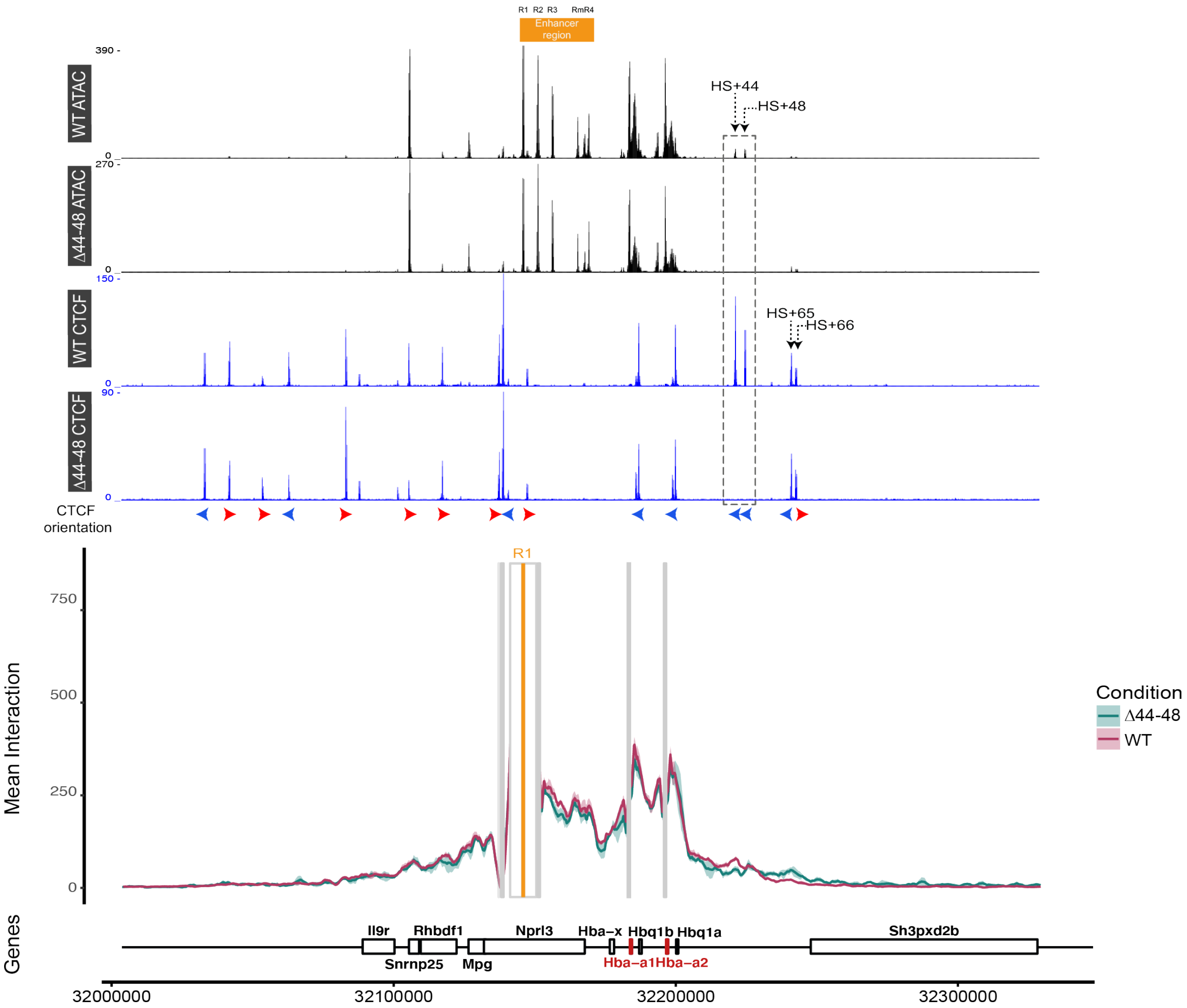


Top tracks show profiles for ATAC-seq and CTCF ChIP-seq in primary erythroid cells isolated from WT (26) and Δ44-48 mice (Ter119+) for the α-globin locus on chromosome 11. Profiles show normalised (RPKM) and averaged data from three biological replicates. The orientation of CTCF motifs is shown under peaks by red (forward) and blue (reverse) arrows. NG Capture-C interaction profiles of the α-globin locus from the viewpoint of R1 (orange), with a 1 kb exclusion zone around the viewpoint, in WT (red) and Δ44-48 (green) Ter119+ primary erythroid cells. The profiles represent normalised and averaged unique interactions from three biological replicates, with the halo representing the standard deviation of a sliding 3 kb window. Grey bars denote other capture points included in this experiment. Genes and genomic position below interaction profiles with the α-globin genes highlighted in red.

**Supplementary Figure 3: Chromatin accessibility, CTCF binding, and quality of NG Capture-C profile in Δθ1 erythroid cells**

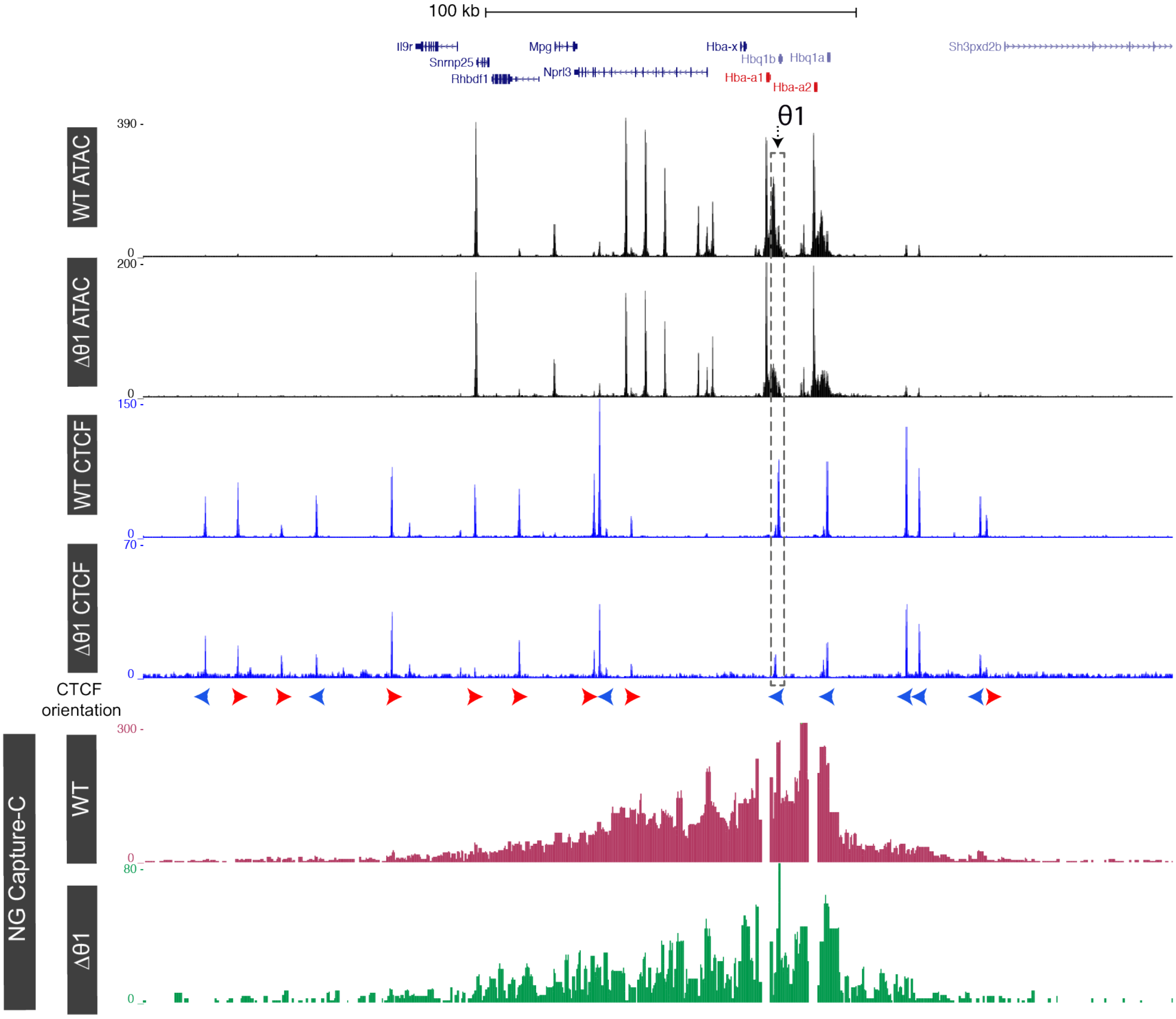


Top tracks show profiles for ATAC-seq and CTCF ChIP-seq in primary erythroid cells isolated from WT (26) and Δθ1 mice (Ter119+) for the α-globin locus on chromosome 11. Profiles show normalised (RPKM) and averaged data from three biological replicates. The orientation of CTCF motifs is shown under peaks by red (forward) and blue (reverse) arrows. NG Capture-C interaction profiles of the α-globin locus from the viewpoints of the α-globin promoters, with a 1 kb exclusion zone around the viewpoints, in WT (red) and Δθ1 (green) Ter119+ primary erythroid cells. The profiles show one representative biological replicate to highlight the low complexity and poor-quality library in Δθ1 erythroid cells, and any differences between the model and WT could not be determined. Gene annotation from Refseq with the α-globin genes highlighted in red.

Preliminary analysis suggests that this model may harbour SNPs in the DpnII sites around the α-globin locus originating from the genetic backgrounds of the mice used to generate the model. The presence of SNPs in DpnII sites impacts the digestion of 3C libraries and drastically alters the complexity of interaction profiles, thus we were unable to generate sufficient interaction profiles from this model.

**Supplementary Table 1: DNA sequences for sgRNAs and ssODNs used for targeting CTCF binding sites at the mouse a-globin locus**

| Target | Guide Sequence (5'-3') | ssODN Sequence (5'-3') |
|---|---|---|
| **θ1** | TGGAACGATGCAGCGCCCCC | CGCTCTGCCCGCTGGCTGAGCTCAAAGACGTCCTGAAACACAAGAGGCGGATCCCGCTGCATCGTTCCAGGATGCCTAGGTGTTCACAGATTCTGGTTCAGCTTTGAGCCCTCTGTTTCCCTGGGCTCCCCCTCC |
| **θ2** | TGAAACACAAGAGGCCGCCA | CTTGCGCTCTGCCCCCTGGCTGAGCTCAAAGATGTCCTGAAACACAAGAGGCGGATCCCGCTGCATCGTTCCAGGATGCCTAGGTGTTCACAGATTCTGGTTCAGCTTTGAGCCCTCTGTTTCCCTGGGCTCCCT |
| **θ2** | GACATCTTTGAGCTCAGCCA | |
| **HS+44** | GAAAGCCAGTGGCGCCACCT | N/A |
| **HS+44** | CCCTGCAGGCCACTATAAGT | N/A |
| **HS+48** | TCCAAGGTCCTCAAGCAGAC | N/A |
| **HS+48** | CGACGAGCACCCCCGTGTGG | N/A |
